# Assessing climate adaptation among Canada lynx (*Lynx canadensis*) populations at the trailing edge

**DOI:** 10.64898/2026.02.04.703807

**Authors:** Tanya M. Lama, Blair P. Bentley, Warren E. Johnson, Lisa M. Komoroske, Stephen Destefano, Jennifer Vashon, Kaela Beauclerc, Jason M. Kamilar, Giulio Formenti, Andrea Batista, Andrew G. Spencer, Olivier Fedrigo, Erich D. Jarvis, Jeff S. Bowman, Paul Wilson, John F. Organ

## Abstract

Species must acclimate, shift their distributions, or adapt in place in response to anthropogenic climate change. Populations at low-latitude trailing edges of species distributions typically experience thermal conditions closest to the upper limit of their thermoregulatory capacity. Landscape and functional genomic approaches provide quantitative measures of risk and adaptive capacity which can inform and prioritize conservation actions. Using low-coverage whole genomes from Canada lynx (*Lynx canadensis*), we characterized population genomic structure and identified putatively adaptive loci using genotype-environment association analyses across the eastern extent of their distribution. We detected genetic breaks across two previously identified biogeographical barriers, the St. Lawrence River and the Strait of Belle Isle, and found relatively high genome-wide diversity in the Maine population at the southern trailing edge, suggesting a reservoir of warm-adapted variation. We identified 759 loci from 329 genes as putatively adaptive, many associated with temperature during warm and dry periods, and functionally enriched in photoreception, circadian entrainment, and temperature regulation. We identified ten putatively adaptive genes linked to epilepsy, presenting candidate genes underlying reports of idiopathic epilepsy in captive populations of closely related lynx species (*L. lynx* and *L. pardinus*). Genetic offset showed lynx in Western Newfoundland, and the Gaspé Peninsula in Quebec are at the greatest risk of maladaptation under future conditions. If gene flow allows, introgression of climate-adapted loci from the trailing-edge may benefit regional populations under future climates. Together, these findings demonstrate the conservation value of locally adapted range-edge populations.

## Introduction

Global climate change will soon surpass habitat destruction as the leading threat to biodiversity (Pereira et al. 2010; Urban 2015), spurring ecological responses across broad geographic scales (Parmesan 2006). Species persistence in a rapidly changing climate relies on spatial redistribution, phenological shifts, acclimation through plasticity, and evolution towards new behavioral and physiological optima (Crozier et al. 2008). Range shifts in response to climate change have been studied extensively across taxa, and described in all directions and at varying rates and magnitudes (MacLeod 2009, Lenoir and Svenning 2015). However, few species are capable of the large-scale dispersal required to redistribute to landscapes that include their complete suite of optimal habitat conditions (Gienapp et al. 2008; Visser 2008; Hanski 2012). For example, species that redistribute to maintain access to one aspect of their habitat requirements (e.g. prey abundance), might simultaneously become discordant with other critical habitat needs. To compound these challenges, redistribution is likely to be hampered by habitat fragmentation in human-dominated landscapes. Thus, it is likely that many species will in part rely upon rapid population-level microevolution to adapt on pace with the unprecedented rate of climate change (Visser 2008).

Genetic adaptation to a changing environment occurs through selection on novel mutations and pre-existing “standing” genetic variation (Barrett and Schluter 2008). Compared to new mutations, adaptation from standing genetic variation is likely to be faster, and standing variants have been “battle-tested” by selection in a variety of different contexts (e.g. under historic conditions, from another part of the species’ range, or introgressed from another species) (Barrett and Schluter 2008). For example, beneficial alleles present among standing variation are immediately available within the population and may exist at higher frequencies than new mutations, reducing their time to fixation (Scheffers et al. 2016). Therefore, evaluating a species’ evolutionary potential, or ability to evolve in response to climate change, considers the richness and distribution of standing genetic variation through allele and genotype frequencies, diversity indices, and measures of population differentiation along with ecological factors (Barrett and Schluter 2008; Bitter et al. 2019).

Forester and Lama (2023) described barriers and opportunities for better integrating quantitative assessments of evolutionary potential into Endangered Species Act (ESA) listing and recovery decisions. The availability of well-annotated reference genomes, such as those publicly available through the Vertebrate Genomes Project (Rhie et al. 2020), is quickly lifting barriers for species of conservation concern (Formenti et al. 2022). Importantly, these genomic resources empower us to describe the spatial and temporal distribution, and functional significance, of adaptive genotypes (Hohenlohe et al. 2021; Theissinger et al. 2023). The distribution of adaptive genotypes is especially important when considering climatic change, as local adaptation – the evolutionary process whereby populations possess traits that confer higher survival and reproduction under local conditions – will likely result in a mismatch between genotypes and future conditions (Capblancq et al. 2020).

Populations that exist at the range-edge of a species distribution are thought to be adapted to the extremes of a species’ fundamental niche (Angert et al. 2020), and the trailing edge of a species distribution presents a natural experiment to test whether adaptation can outpace local extinctions (Hampe and Petit 2005). For example, at the southern trailing edge, species’ responses to climate change should be more apparent, where populations are small, suitable habitat tends to be patchy, reproductive success is low, and mortality is high (Gaston 2009).

Trailing edge populations may also harbor unique adaptive variants that are beneficial under future conditions (Hampe and Petit 2005). Therefore, range edge populations may serve a functionally important role in safeguarding potentially climate-adapted variation. Genomic forecasting allows us to identify populations at the greatest risk of climate maladaptation, when the connection between a fitness optimum and its underlying genotype is disrupted in locally adapted populations (Fitzpatrick and Keller 2015). Such predictive methods use population genomic data to assess future mismatches in locally adapted gene-environment associations (Capblancq et al. 2020), with higher degrees of mismatch (i.e. genomic offset; Fitzpatrick and Keller 2015) indicating higher susceptibility to the impacts of climate change. Hence, genomic forecasting presents a valuable tool for identifying populations at the greatest risk of local extinction (Bay et al. 2018; Ruegg et al. 2018), protecting unique and climate-adapted genotypes, and informing assisted gene flow between populations (Aitken and Whitlock 2013; Steane et al. 2014; Aitken and Bemmels 2016).

Canada lynx (*Lynx canadensis*) are cold-adapted specialists common across the boreal forests of Canada and Alaska, but populations at the southern edge of their range are protected under both the U.S. Endangered Species Act and Canadian Species at Risk Act (Koehler and Aubry 1994; USFWS 2000). The northeastern United States is projected to warm faster than many other global regions (Karmalkar and Bradley 2017) presenting climate-related threats to lynx at the trailing-edge including rising temperatures, reduced snowpack, and shorter periods of snow persistence (King et al. 2020). Although long-term warming is evident, winter conditions over the past ∼30 years show substantial temporal and spatial variability in snowfall and temperature, including years with exceptionally low (e.g., 2004, 2010) or high snow accumulation (e.g., 2008, 2019). Notably, parts of northern Maine continue to receive consistently high snowfall, suggesting that localized climate refugia may persist despite regional trends. The trailing edge of the Canada lynx distribution in the contiguous U.S. and southern Canada has contracted markedly (>175 km) over the past century (Koen et al. 2014; Thornton and Murray 2024), though the exact magnitude remains uncertain (Ivan et al. 2024). Potential persistence of populations in this region is further challenged by major geographic barriers that limit gene flow between the trailing edge and core populations, including the St. Lawrence River and the Strait of Belle Isle (Koen et al. 2014; Prentice et al. 2017). These barriers may also restrict the movement of potentially warm-adapted alleles from edge populations into the core, limiting the species’ evolutionary response to climate change.

Here, we assessed the risk of maladaptation to future climates across the eastern distribution of Canada lynx, including trailing edge populations in Maine and eastern Canada. We (1) estimated gene flow occurring between populations and the impacts of geographic barriers; (2) estimated evolutionary potential and inbreeding; (3) identified putatively adaptive loci and their functions; and (4) estimated genomic offsets for each population under future climates. Together, we present the first assessment of evolutionary potential for Canada lynx, providing an important baseline for the recovery and monitoring of range-edge populations in the contiguous United States.

## Materials and Methods

### Canada lynx distribution in Maine and eastern Canada

Canada lynx are common in the boreal forests of Alaska and Canada, and their distribution coincides with that of their primary prey, the snowshoe hare (*Lepus americanus*). Lynx in the contiguous United States are at the southern trailing edge of the species distribution and were once found in at least 14 northern states in the Cascade and Northern Rocky Mountains, Western Great Lakes, and the Northeast (USFWS 2000). In the Northeast, their distribution is limited to the upper extent of the Northern-Appalachian ecoregion, where regenerating conifer and mixed forests support abundant snowshoe hare populations, and winter conditions confer a seasonal advantage over generalist mesocarnivores (Hoving et al. 2005; Carroll 2007). Our study area encompasses lynx in Maine and adjacent Canadian provinces (Quebec [north and south of the St. Lawrence River (SLR)], New Brunswick, Labrador, and the island of Newfoundland (Fig. 1). Population structure in this region has been characterized in prior microsatellite-based analyses (Koen et al. 2014; Prentice et al. 2017).

**Figure 1.**
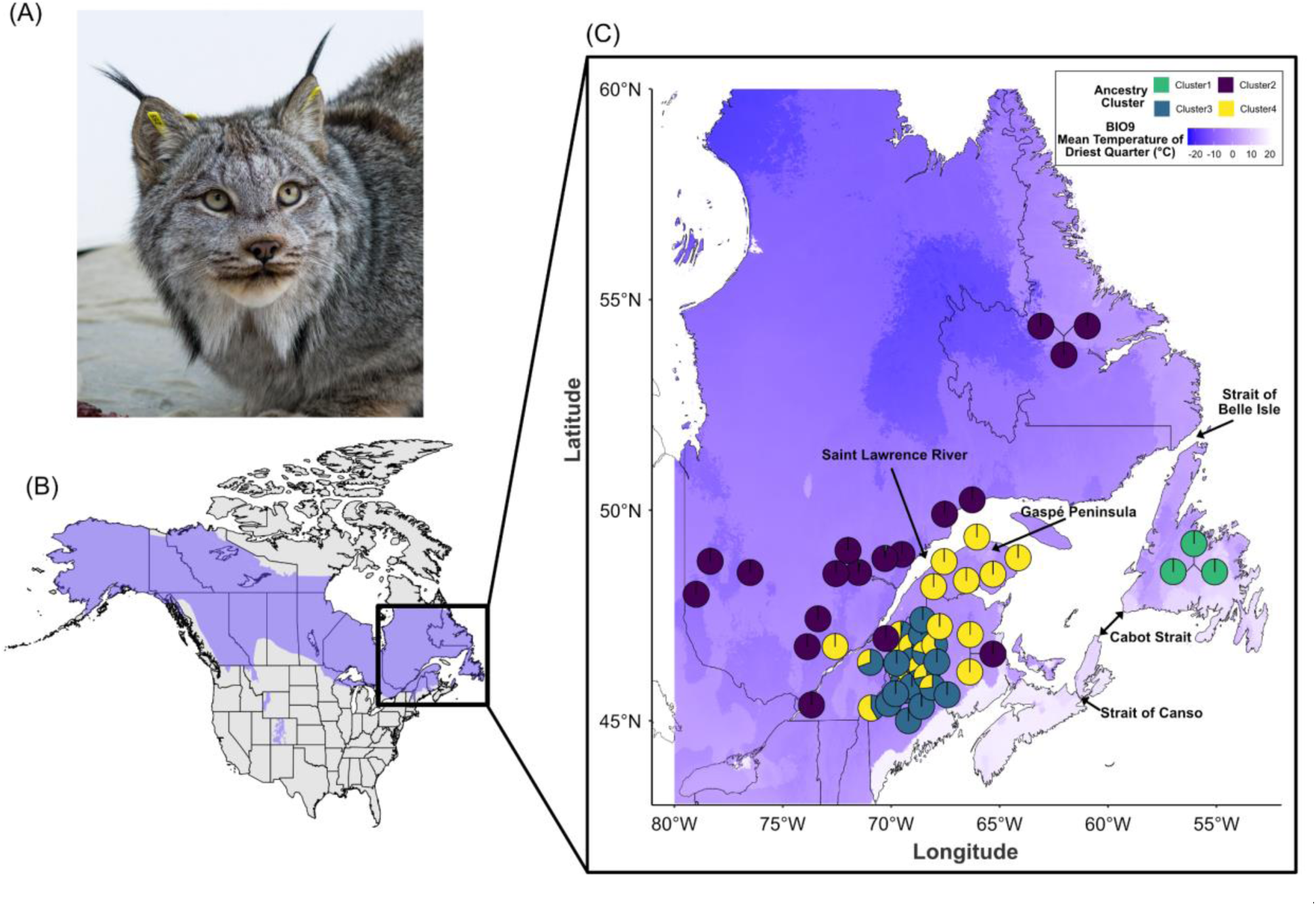
(A) Canada lynx (*Lynx canadensis*; photograph by Bill Byrne). (B) Canada lynx are distributed across North America (highlighted purple region), including the northern Appalachian-Acadian ecoregion throughout Maine and eastern Canada (boxed region). (C) Fifty-seven Canada lynx samples were collected from across this region between 2012-2018. Quebec province samples are spatially referenced to the centroid of harvesting units called Unites de Gestion des Animaux a Fourrure (UGAF). Samples from Labrador, Newfoundland, and New Brunswick are plotted as the centroid of the respective province. Maine samples are spatially referenced to roadkills and incidental capture events during the fur-trapping season. Map shading represents the mean temperature of the driest quarter (BIO9) across the sampled region. Pies represent assignment of each individual to four genetic clusters inferred from admixture analyses using whole-genome genotype-likelihoods. Clusters identified through Admixture analysis (see *Results*) broadly represented individuals from Quebec north of the St. Lawrence River & Labrador (purple; NSLR), Quebec south of the St. Lawrence River and New Brunswick (yellow; SSLR), Maine (blue), and Newfoundland (green).

### Sample collection

Frozen and desiccated tissue samples (muscle, N=26) from Maine were acquired from the Maine Department of Inland Fisheries and Wildlife (MDIFW). Samples from 2016-2018 were collected from live animals incidentally captured during the Maine fur-trapping season and released, or collected post-mortem from roadkill incidents. Samples acquired from Canadian provinces [total N=32; Quebec (N=23), New Brunswick (N=3), Labrador (N=3), and Newfoundland (N=3)] from 2009-2016 are the same as those described in Prentice et al. 2017, albeit with smaller sample sizes per location (see Prentice et al. 2017 for detailed sample descriptions).

### DNA extraction and sequencing

Genomic DNA was extracted from tissue using the PureLink™ Genomic DNA Mini Kit (Invitrogen #K182002). DNA quality and quantity was evaluated on a Fragment Analyzer (Agilent Technologies, Santa Clara, CA) using standard-(DNF-467-0500) and high-sensitivity (DNF-468-0500) kits. Only samples that showed clear high molecular weight bands and limited smearing were retained for library preparation. DNA yield and degradation varied among samples, but usable DNA was extracted from 57 samples. A separate barcoded whole-genome library was prepared for each individual according to the protocol optimized for high-throughput, low-cost genome generation by Therkildsen and Palumbi (2017). The final concentration of input DNA ranged from 1.3 - 5.3 ng/mL for each library. Due to international permit limitation, samples were sequenced over two independent runs. Equimolar amounts of library were pooled and sequenced on one lane of paired-end 125-bp reads on an Illumina NovaSeq S4 flow cell at Novogene Corporation (Sacramento, CA, USA), and one lane of paired-end 125-bp reads on an Illumina NovaSeq S1 flow cell at The Centre for Applied Genomics (Toronto, ON, Canada). Scripts for all downstream analyses are publicly available on GitHub: https://github.com/bpbentley/lynx_wgr

### Genotype likelihoods

Raw data were screened for quality with FastQC (v0.12.1; Andrews et al. 2012), with residual adapter sequences, low quality base calls, and small read lengths trimmed using the BBduk function of BBMap (v39.26; Bushnell 2014). Trimmed reads were aligned to the Canada lynx reference genome (GCA_007474595.2; Rhie et al. 2020) using bwa-mem (v0.7.19; Li 2013) with default parameters, before PCR duplicates were removed using picard tools (v3.3.0; Broad Institute 2019). Downstream mapping and alignment statistics for quality control were calculated using samtools (v1.21; Li et al. 2009) and bedtools (v2.31.1; Quinlan and Hall 2010). We used the Analysis of Next Generation Sequencing Data (ANGSD; v0.935; Korneliussen et al. 2014) software to generate genotype likelihoods directly from the final alignment files.

Genotype likelihoods were used in place of hard genotype calls due to relatively modest depth coverage (4-15×) of the sequenced individuals. Likelihoods were generated in ANGSD using the GATK algorithm (-GL 2), with SNP p-value of 2e^-6^, a minimum minor allele frequency of 0.05, minimum base quality of 20, minimum mapping quality of 30, and present in at least 30 individuals.

### Population genomic structure and diversity

For population structure, we conducted a principal component analysis (PCA) directly on the beagle file output of genotype likelihoods from ANGSD using PCAngsd (Meisner and Albrechtsen 2018), and visualized outputs with ggpubr (Kassambara 2025) in R version 4.5.0 (R Core Team 2025). Admixture proportions were generated from the genotype likelihood data using NGSadmix (Skotte et al. 2013), with 10 replicates of each cluster assignment (K=2-6). Following admixture analyses, individuals were partitioned into their respective clusters (K=4) for downstream analyses, with individuals assigned >30% secondary ancestry considered ‘admixed’. ANGSD was subsequently applied to generate unfolded site allelic frequencies (SAF;-doSaf) independently for each cluster, followed by folded pairwise 2D site frequency spectrum (realSFS) comparisons between all clusters using unfolded SAFs. Finally, weighted pairwise F_ST_ between each cluster (not incorporating admixed individuals) was calculated using the ‘fst’ function implemented through the ANGSD realSFS subroutine.

Heterozygosity (Watterson’s theta) and runs of homozygosity (ROHs) for each individual were calculated using the Bayesian framework method implemented through ROHan (Renaud et al. 2019). The ROHan routine was estimated on individual alignment files, with a Ts/Tv ratio of 2.4 (calculated from the VCF of hard genotype calls using vcftools; see *Genotyping through base calls*), with the default values of 1 megabase window size and a rohmu of 2e^-5^. The ROHan analyses were run excluding sex chromosomes and unplaced scaffolds, with the remaining sequence covering eighteen chromosomes (94.8% of the total reference assembly). ROHan has been demonstrated to perform relatively well on low-to moderate-coverage sequencing depths (Taylor et al. 2026). Heterozygosity estimates for low-coverage data were additionally validated using ANGSD; by generating folded SAFs on autosomes from each alignment file, filtering for excessive mismatches (-C 50), and for base and mapping quality of less than 20 and 30, respectively. The realSFS subroutine was then used on the individual SAF files, and heterozygosity was estimated using R.

### Genotyping through base calls

Genotype-environment associations identify adaptive loci using hard-called genotype inputs, rather than genotype likelihoods. Resequenced genomes with <4× coverage were removed from hard-call genotyping, leaving a total of 55 individuals. Variants were identified from alignment files described above (see *Genotype likelihoods*) using the Genome Analysis Toolkit (GATK v4.5.0.0) with the best practices workflow. Briefly, variants were filtered to retain only biallelic SNPs with a mapping quality of >40, a total locus depth between 15-1500× across all samples, and a minimum individual depth of >4× per locus. SNPs were also filtered to remove loci missing in >30% of samples and with a minor allele frequency of <0.05 (full details of filtering steps are available on GitHub).

### Estimation of effective migration surfaces

We leveraged the hard-called SNP set to estimate effective migration surfaces (EEMS) across the studied region (Petkova et al. 2016). EEMS predicts the connectivity of populations across geographic scales by assessing the decay of genetic similarity between sampling locations under a null hypothesis of isolation-by-distance. A dissimilarity matrix of genotypes was created using the bed2diffs function in EEMS from the VCF file converted to PLINK format.

Geographic distance was computed internally through the EEMS analysis, using the GPS coordinates of the sampling locations, and an outer boundary limit generated from the Canada lynx distribution across the region. EEMS was run independently three times, with 200 demes, 100,000 MCMC iterations, and 1,000 burn-in chains. The chain with the lowest log likelihood was selected, and visualized using the ‘eems.plots’ function of the rEEMSplots package for R included in the EEMS program.

### Climate data

Bioclimatic variables derived from monthly temperature and precipitation data served as a proxy to describe climatic conditions relevant to Canada lynx. Using GPS coordinates associated with each sample collection location, raster data for 19 bioclimatic variables (BIO1 to BIO19) were extracted at 30-second arc spatial resolution from the WorldClim version 2.1 database (Fick and Hijmans 2017) for the period between 1970 and 2000, hereafter described as the “current” climate. Raster data for all “future” bioclimatic variables (BIO1 to BIO19) were also extracted from Coupled Model Intercomparison Project Phase 6 (CMIP6) projections at 30-second arcs. CMIP6 data were selected using the “Middle of the Road” Shared Socioeconomic Pathway (SSP2-4.5; Thomson et al. 2011) projection for four time periods: (1) 2030 [2021-2040]; (2) 2050 [2041-2060]; (3) 2070 [2061-2080] and (4) 2090 [2081-2100] (Eyring et al. 2016). The “Middle of the Road” SSP2-4.5 narrative broadly describes medium challenges to mitigation and adaptation to global climate change, in which “social, economic, and technological trends do not shift markedly from historical patterns” (van Vuuren et al. 2017). In other words, we selected the “business as usual” scenario for future climate projections (Hausfather and Peters 2020).

We initially calculated correlations between bioclimatic variables (Fig. S1) and removed variables that showed high pair-wise correlations using the ‘findCorrelations’ function of the caret R package (Kuhn 2008), with a stringent correlation cutoff of 0.8. This correlation filter resulted in eight bioclimatic variables being retained. However, due to *a priori* biological interests, we manually switched BIO7 (Temperature Annual Range) with BIO19 (Precipitation of Coldest Quarter) for downstream use. These eight variables were also applied for all future scenarios, with the remaining bioclimatic variables excluded from all analyses.

### Outlier tests and genotype-environment association (GEA) analyses

Three independent methods were used to identify putatively adaptive loci. PLINK (v1.90b7.7; Purcell et al. 2007) was used to remove loci in linkage disequilibrium (LD;--indep-pairwise 50 10 0.2) from the filtered hard-called SNP. An initial outlier test was performed on the LD-pruned VCF using the pcadapt package in R and an optimal K value identified by scree plotting (Fig. S2). The resultant p-values were corrected for multiple comparisons using the qvalue package in R and loci with an adjusted p-value of <0.05 were identified as outliers (Fig. S3).

Two genotype-environment association (GEA) analyses were used to complement the outlier analysis, and to associate specific bioclimate variables with identified loci. As GEA analyses are sensitive to missing data, we initially imputed missing genotypes in our dataset using the ‘impute’ function of the LEA package (Frichot and François 2015), leveraging the results of a sparse Non-Negative Matrix Factorization analysis (sNFM; Frichot et al. 2014) with the ‘mode’ parameter and K=4. With the imputed dataset, a latent factor mixed model (LFMM, Caye et al. 2019) procedure was conducted using the eight retained bioclimatic variables scaled with one latent factor (K=4). As with the imputation, our latent factor was defined *a priori* by simple non matrix factorization (sNMF) and PCA, and p-values were corrected for multiple comparisons using qvalue. As LFMM conducts individual univariate analyses on each bioclimatic variable, unique loci were identified, and an upset plot (UpSetR; Conway et al. 2017) was used to visualize overlapping loci between BioClim variables (Fig. S4).

We also used a multivariate redundancy analysis (RDA) following methods by Forester et al. (2018). The RDA was applied to the same imputed genotype data and the same scaled bioclimatic variables described for the LFMM analysis, and conducted using the rda function of the ‘vegan’ package (Oksanen et al. 2025). A biplot was created to visualize contributions of specific bioclimatic variables on genetic variation, and candidate SNPs were identified on each axis as those more than three standard deviations away from the mean (i.e., two-tailed p-value < 0.0027). We compared the list of outliers found using the three independent approaches and retained loci identified as outliers using at least two of the methods (see Supplementary Information). This consensus set of outliers have been described as an “adaptively enriched genetic space” (Steane et al. 2014) encompassing putative adaptive genetic variation strongly associated with current climate.

### Functional annotation of adaptive loci and gene set enrichment analysis

The resultant set of putatively adaptive genes was annotated using the NCBI RefSeq Canada lynx reference genome annotation (Rhie et al. 2020; Annotation Release 102). The biological function of putatively adaptive genes was described using ‘gprofiler2’ in R (Kolberg et al. 2020). The g:Profiler program is a gene set enrichment analysis (GSEA) that sorts genes into functional categories (molecular function, biological process, cellular component) and assigns them to pathways, and nested gene families annotated with gene ontology (GO) terms (Raudvere et al. 2019). The g:Profiler GSEA was applied using the *Homo sapiens* gene set as the reference set, and a Benjamini-Hochberg FDR corrected p-value of less than 0.05. Gene sets were also screened for enrichment with functional annotations from DAVID (Huang et al. 2009) to identify KEGG GO pathways associated with our candidate gene set. Finally, genetic load was assessed using SnpEff and SnpSift version LGPLv3 (Cingolani et al. 2012a, Cingolani et al. 2012b) to identify impactful variants. Variants were assigned to annotated genes if they were within or adjacent (5kb upstream and downstream) to genes with known functions. The outputs of SnpEff were then used to determine the function and severity of each identified variant (e.g., missense mutation).

### Generalized dissimilarity models and genomic offset under future climates

We used generalized dissimilarity modeling (GDM), a non-linear method to quantify genetic offset for putative climate-adaptive genetic variation identified through outlier analysis and GEAs. GDM is a distance-based method that fits a single regression model to describe the relationship between spatial and environmental variables and genetic variation, and partitions variance in the model between environmental variables and spatiotemporality. This approach estimates the role of each variable in structuring genetic variation (i.e. testing isolation by environment) as a “turnover rate” of allelic compositional change along environmental gradients (Fitzpatrick and Keller 2015). Spatial data, based on GPS coordinates for each sample, and environmental variables (see Climate data) were used as model predictors. GDM was implemented in the gdm package for R (Fitzpatrick et al. 2025) using Euclidean distance to account for the genetic and geographic distances (geo=TRUE) among all individuals. We used the ‘gdm’ function to identify associations among genetic variation, environmental, and spatial variables under current climate conditions. Genetic offsets under the future climate scenarios (2030, 2050, 2070, and 2090) were then estimated, first using the ‘predict’ function with future climate data, then by estimating the Euclidean distance between current and future associations weighted by variable importance for each grid cell on the landscape. A similar procedure was followed for GDM results using the ‘gdmtransform’ and GDM ‘predict’ functions with principal components to identify and map current genotype-environment relationships onto the landscape. The GDM ‘predict’ function was used with current and future climate data in raster format (time=TRUE) to estimate genetic offset in a single step.

## Results

### Canada lynx in Maine have high genomic diversity despite biogeographic barriers

Using principal-component and admixture analyses from genotype likelihoods, our results show distinct population structure across the south-eastern trailing edge of the Canada lynx range (Fig. 2). Consistent with previous findings, we demonstrate clear separation of the isolated Newfoundland population from all other sampled areas. Our results also show a genetic break between samples collected from Quebec North of the St. Lawrence River and Labrador compared to those collected from South of the River, suggesting that this geographic feature impedes gene flow between lynx populations. Interestingly, we also found evidence that samples collected from Maine were genetically distinct from those collected in Quebec SSLR and New Brunswick, suggesting this area may harbor genomic reservoirs of diversity at the distribution’s trailing edge. When considering admixture with four clusters (K=4; Fig. 2C), individuals generally separated out into Newfoundland (hereafter ‘cluster 1’), Quebec NSLR and Labrador (’cluster 2’), Maine (’cluster 3’), or Quebec SSLR and New Brunswick (’cluster 4)’. In addition, eight individuals from Maine, and one from Quebec SSLR were considered ‘admixed,’ showing >30% of secondary ancestry. All admixed samples showed a combination of ancestry from both cluster 3 and cluster 4, suggesting gene flow between these areas. As expected from previous research and island theory, individuals from Newfoundland showed very low estimates of genomic diversity (Fig. 2B), and very high proportions of their genome in ROH (Fig. 2D), suggesting high levels of inbreeding. In contrast, individuals from Maine showed the highest genomic diversity, and lowest rates of inbreeding, suggesting high standing variation.

**Figure 2.**
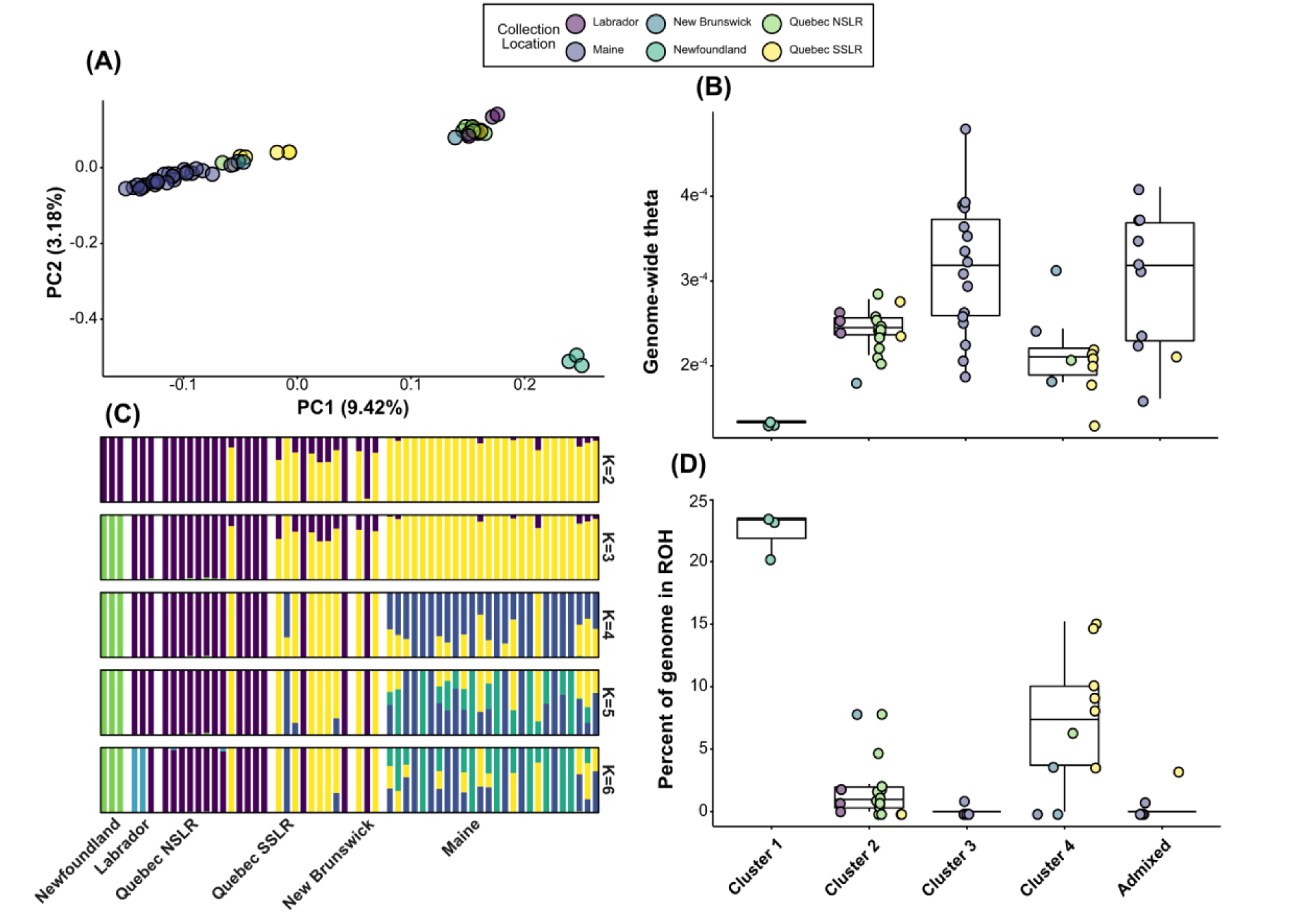
Genetic structure, diversity and inbreeding. (A) Principle component analysis of genotype likelihoods from fifty-seven individuals across the southeast trailing edge of the Canada lynx distribution. (B) Genome-wide diversity (theta) of all individuals, separated by cluster assignment with K=4. (C) Admixture analyses for K=2 through K=6. (D) Overall percentage of individual genomes that were categorized as being in runs of homozygosity (ROH) with a minimum length of 1megabase as the default window size parameter for ROHan.

### Limited gene flow across the St. Lawrence River and the Strait of Belle Isle

Outputs from weighted pairwise F_ST_ comparisons and estimations of effective migration surfaces (EEMS) were consistent with results from the principal component and admixture analyses. Pairwise-F_ST_ was highest between the Newfoundland and Maine clusters (F_ST_ = 0.236), and the Newfoundland and Quebec SSLR/New Brunswick clusters (F_ST_ = 0.233; Fig. 3A). In contrast, pairwise F_ST_ was lowest between Maine and Quebec SSLR/New Brunswick (F_ST_ = 0.052), even when excluding the admixed individuals, further suggesting relatively higher rates of gene flow between these areas. The pairwise F_ST_ between Newfoundland and Labrador/Quebec NSLR suggested restricted gene flow (F_ST_ = 0.168), but is slightly lower than the other comparisons with the island.

**Figure 3.**
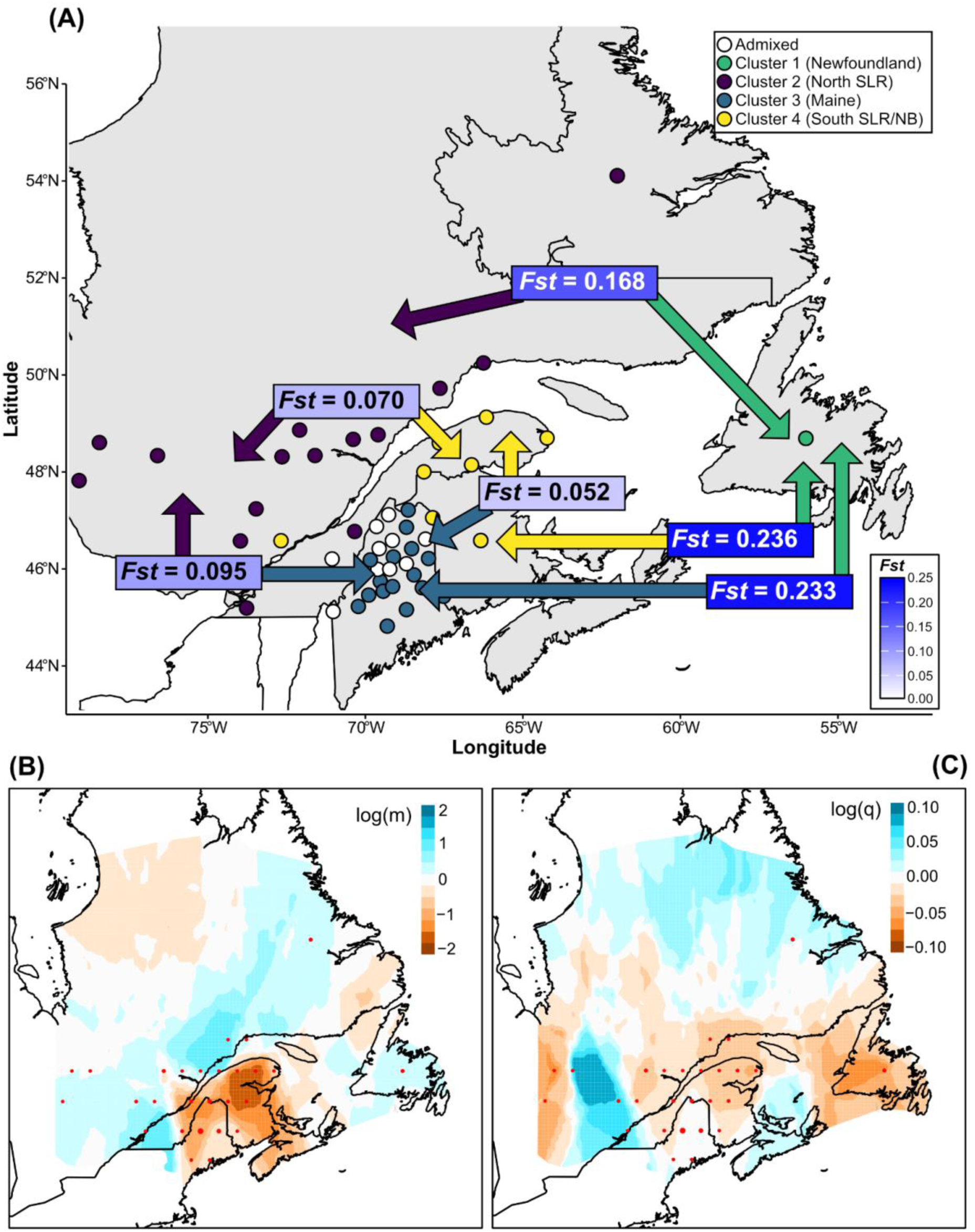
(A) Weighted pairwise F_ST_ generated for all genotype likelihoods between clusters identified through NGSadmix (K=4). Admixed individuals (white; secondary ancestry >30%) were excluded from pairwise comparisons. Colored arrows represent the pairwise comparison, and F_ST_ boxes are colored on a gradient from 0 to 0.25 (B) Estimation of Effective Migration Surface (EEMS) contours illustrate relative migration. Each circle (red) is a deme consisting of one or more individuals. Color scale indicates relative migration across the study area with orange contours indicating low levels of migration and blue contours indicating high levels of migration. (C) EEMS contours illustrate the effective diversity rate within each deme (red). Color scale indicates genome-wide diversity across the study area with orange contours indicating lower than expected diversity and blue contours indicating higher than expected diversity.

While the pairwise F_ST_ analysis requires supervised clustering of individuals, the EEMS analysis relies on matrices of genetic and geographical distances, and is thereby unbiased in the outputs. Our analyses showed that migration rates are relatively low across the St. Lawrence River (log_m_ = −1), and between Newfoundland and Labrador (log_m_ = −0.05; Fig. 3B).

Surprisingly, the EEMS analysis also showed low levels of migration between Maine, Quebec SSLR, and New Brunswick (log_m_ = −1.5 to −2), despite previous analyses suggesting that gene flow is high in this area and admixture is common. In addition to estimating migration, the EEMS analysis demonstrated regions of lower-than-expected heterozygosity. Consistent with previous results, Newfoundland shows a strong signal of reduced genomic diversity (log_q_ = - 0.07; Fig. 3B).

### Concordant identification of putatively adaptive loci

From a total panel of 92,047 LD-pruned genotyped SNPs across our samples, 759 (0.82% of all LD-pruned SNPs) loci were identified as putatively adaptive using at least two of the three independent outlier detection methods (PCAdapt, LFMM, and RDA). These analyses detect loci showing allele frequency patterns that are either outliers with respect to population structure (PCAdapt) or significantly correlated with environmental gradients after accounting for population structure (LFMM and RDA; Fig. S8). The least conservative method, PCAdapt, which does not incorporate climatic data, identified 3,322 SNPs as outliers, while the most conservative LFMM method detected 843 SNPs associated with the BioClim variables, with the majority linked to BIO10 (Fig. S4). The highest number of loci overlapped between the PCAdapt and RDA analyses. From the final set of 759 putatively adaptive SNPs, 379 were linked to annotated genes by SnpEff, which accounts for 2.91% of genes in the NCBI *Lynx canadensis* annotation (Release v102).

### Putatively adaptive SNPs are functionally enriched in sensory perception, thermoregulation, and circadian entrainment

A total of 292 of the 379 genes (77.04%) were assigned to gene ontology terms by g:Profiler and DAVID using the *Homo sapiens* gene set. Screening the gene ontology terms of all the putatively adaptive loci revealed many of these genes were associated with visual perception and photoreception (CEP290, CIB2, CLN8, COL2A1, LCA5, OLFM3, OPN5, PCDH15, SLC7A14); temperature regulation, cold-induced thermogenesis, environmental stimulus, and homeothermy (ARNT2, DBH, NDUFA12, NTSR1, PLCL1); response to heat stress (NUP35, NUP54, PIRT); and circadian rhythm and entrainment pathways (EGFR, NPAS2, OPN5, PLCB4). We additionally detected several genes that have been previously associated with epilepsy and/or other neural conditions that promote seizures (GABRA1, DSCAML1, CLN8, HEXB, KCNH1, KCNMA1, PRICKLE2, CNTNAP2).

Of the 759 putatively adaptive SNPs, none were annotated as high impact by SnpEff, however, seven were annotated as moderately impactful, and a further ten were classified as low impact (Table S2). The moderate impact variants were associated with five genes: CC1H1orf174 (aka CCDC174), LCA5, NDUFA12, and two unannotated genes that are putatively related to olfaction and T-Cell reception. Annotations suggested that in all seven moderate impact variants, the change resulted in a missense substitution (non-conservative amino acid change), potentially impacting the protein function. Variants annotated as low impact were associated with seven genes: DSCAML1, GABRA1, KCNN1, KRT84, PLXND1, PXDN, and one unannotated gene with similarity to glycolipid transfer protein. All low impact variants were predicted to result in synonymous substitutions that did not alter amino acid sequences.

### Enrichment in developmental and neurogenesis-related pathways

Exploring gene set enrichment in the putatively adaptive loci showed, 11, and 16 enriched gene ontology pathways for biological process (BP, n=118), molecular function (MF, n=11), and cellular component (CC, n=16) (Fig. 4A). The five most significantly enriched BP assigned were nervous system development (GO:0007399), system development (GO:0048731), neurogenesis (GO:0022008), multicellular organism development (GO:0007275), and generation of neurons (GO:0048699) (Fig. 4B; Table S3). For MP, the five most significant terms were transmembrane receptor protein phosphatase activity (GO:0005001), transmembrane receptor protein tyrosine phosphatase activity (GO:0019198), protein binding (GO:0005515), phosphoprotein phosphatase activity (GO:0004721), and protein tyrosine phosphatase activity (GO:0004725) (Fig. 4C; Table S4). Finally, for CC, the most enriched terms were synapse (GO:0045202), cell junction (GO:0030054), cell projection (GO:0042995), plasma membrane bounded cell projection (GO:0120025), and cytoplasm (GO:0005737) (Fig. 4D; Table S5).

**Figure 4.**
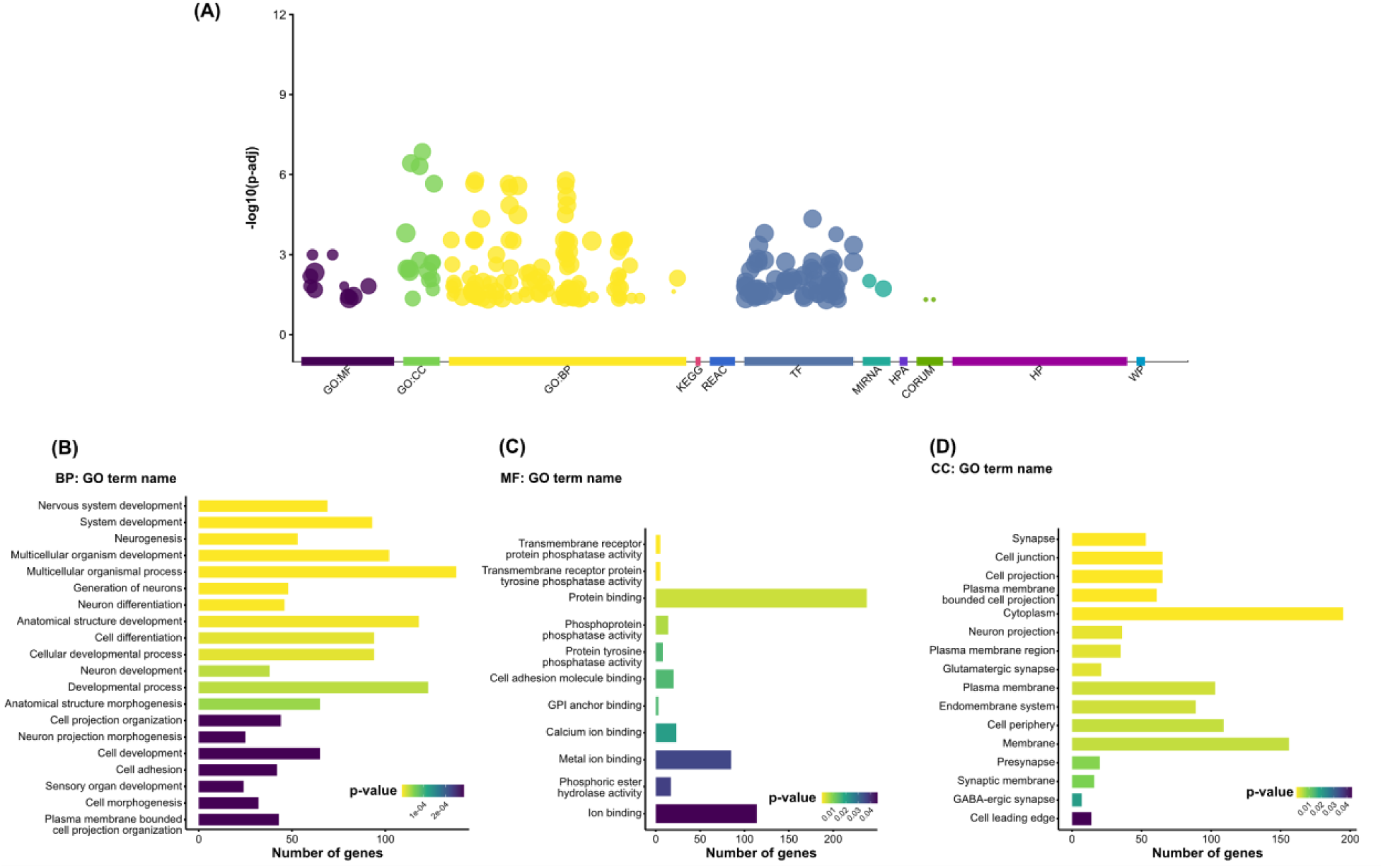
Outputs from the g:Profiler GOSt gene set enrichment analysis on the putatively adaptive set of loci identified across the eastern distribution of Canada lynx (*Lynx canadensis*). (A) Total number and significance of all terms that were identified as enriched. The remaining three panels show the top 20 (or less) most significant enriched terms and total number of genes associated with each for (B) Biological Process [BP]; (C) Molecular Function [MF]; and (D) Cellular Component [CC].

### Genomic offset suggests populations in western Newfoundland and the Gaspé Peninsula are at risk of future maladaptation

Geographic and environmental distances for the variables considered explain 70.73% of allelic variation (deviance) in the generalized dissimilarity model (GDM). GDM-ranked variable-importance identified mean temperature of the driest quarter (BIO9), geographic distance, and isothermality as top predictors followed by variables of low importance (Fig. 5A). Allelic composition changed sharply between −5°C and 0°C for mean temperature during the driest quarter, whereas changes along geographic distance and other environmental variables (e.g. isothermality) occur gradually or with modest step-changes, if at all (Fig. 5B).

**Figure 5.**
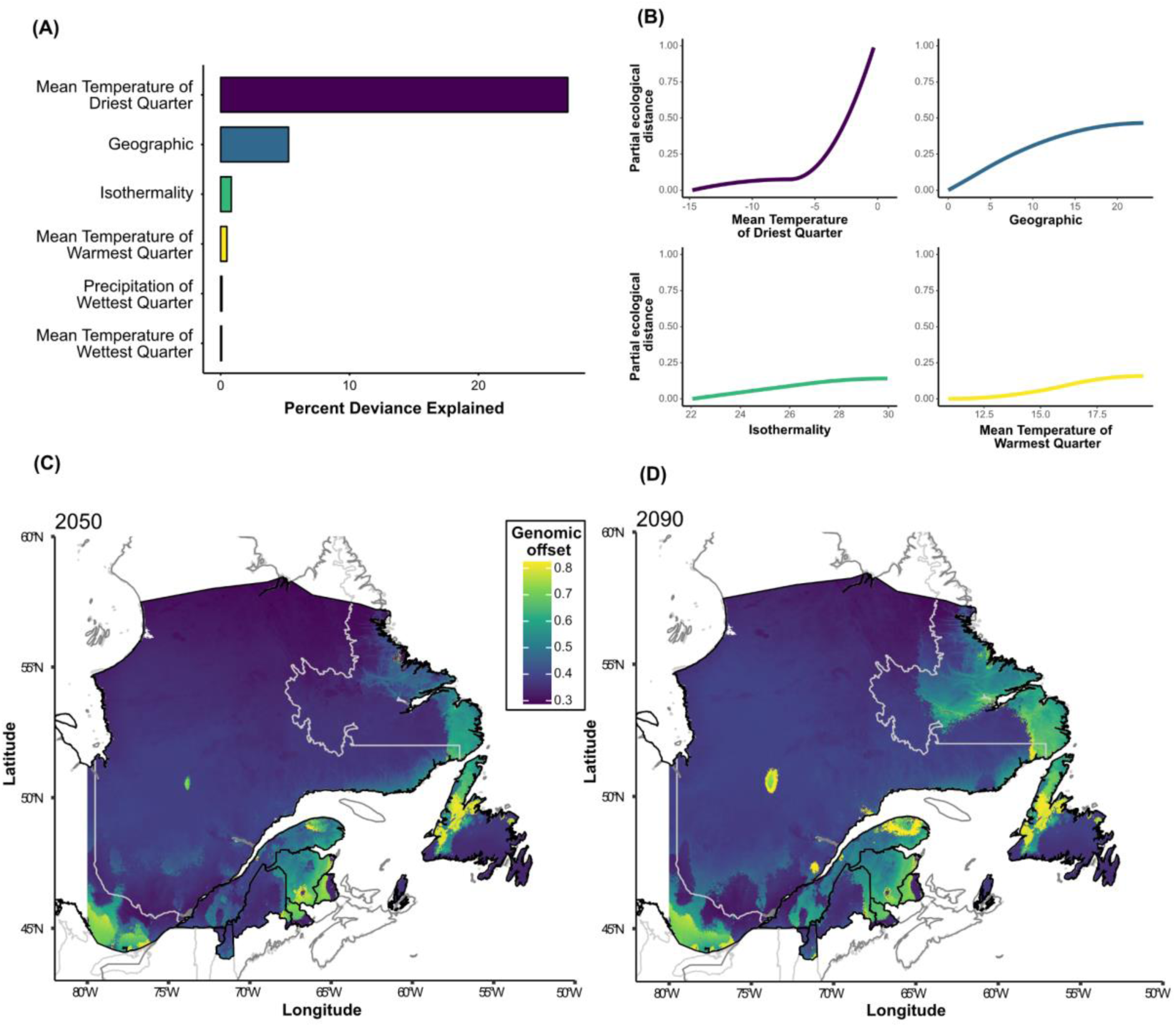
Generalized dissimilarity models (GDM) for predicting genomic offset under future climates. (A) Percentage of model deviance explained by each of the five BioClim variables retained by the GDM and geographic distance. (B) GDM-fitted I-splines for each of the BioClim covariates and geographical distance. The maximum height of each curve shows the total amount of allele frequency turnover associated with that variable. Slope function shape indicates the rate of change in allele frequencies over the variable gradient. (C) Genetic offset for 2050 and (D) 2090 determined by generalized dissimilarity modeling was mapped as the strength of discordance between current and future genotype-environment associations using the identified putatively adaptive loci. Greater discordance (yellow) suggests that stronger allelic shifts will be required for populations to maintain current genotype-environment associations under future conditions.

Mapped projections of GDM results onto the landscape show that peripheral lynx populations in Maine, Labrador, and on the eastern side of Newfoundland, as well as central- and northern-Quebec show the smallest genomic offset (the expected mismatch between current genotype-environment associations and those predicted under future environmental conditions), or vulnerability, in both the 2050 (Fig. 5C) and 2090 (Fig. 5D) predictions. The areas that showed the highest genomic offset were also consistent between the two predicted scenarios, albeit with stronger impacts towards the end of the century. In particular, our projections suggest that Canada lynx in the western half of Newfoundland and areas on the Gaspé Peninsula are at the greatest risk under future climate scenarios. Coastal regions of Labrador, and New Brunswick also showed signs of genomic vulnerability under future conditions.

## Discussion

Urban, Elphick, and Bolnick (2026) illustrated that conservation strategies rarely account for evolutionary potential, and assessments of adaptability have been incorporated into only 4% of federal recovery plans for threatened and endangered species in the US over the past ten years. However, conservation benefits from an evolutionary perspective. Genomic approaches provide quantitative assessments which relate a population’s current condition to the past through neutral variation, and resilience to future challenges through adaptive variation (Fischman et al. 2023). This framework ensures that resources for conservation are efficiently allocated to the protection and restoration of populations with the greatest chance of future persistence. In this study, we provide the first genome-wide assessment of evolutionary potential and maladaptation to future conditions for Canada lynx (*L. canadensis*). Our findings show limited evidence of contemporary gene flow across the St. Lawrence River and the Strait of Belle Isle, and identify the low-latitude trailing-edge population in Maine as a distinct genetic cluster. We show high genome-wide diversity and low evidence of inbreeding within the Maine population, suggesting that abundant standing variation can power evolutionary potential, even on the periphery of the species’ distribution. Genetic vulnerability is also relatively low in Maine, and much of the central core distribution included in our analyses, yet relatively high in Newfoundland, the Gaspé Peninsula, and parts of Labrador. Together, these results present a compelling exception to the central-marginal hypothesis, which posits that peripheral populations exhibit lower genetic diversity because of isolation and population-size effects (Kennedy et al. 2020). Our findings suggest that the southeastern trailing-edge population of Canada lynx in Maine may harbor a reservoir of climate-adapted variation. Functional annotation of putatively adaptive genes suggest that circadian entrainment and photoreception are associated with current conditions, as are genes related to thermoregulation. If gene flow allows, the adaptive alleles in Maine’s lynx population could be introgressed into regional populations to facilitate adaptation to the rapidly changing climate. Future research can further inform policy and practice by assessing the ecological factors shaping high diversity in the Maine population.

### Gene flow, evolutionary potential, and the trailing-edge

In concordance with previous studies on gene flow across Canada lynx populations (Row et al. 2012; Koen et al. 2014; Prentice et al. 2017), our results show clear genetic breaks across the St. Lawrence River and Strait of Belle Isle with implications for peripheral populations in Maine and Newfoundland. The genetic break across the St. Lawrence River supports the hypothesis that this geographic feature forms a barrier to gene flow. Our results add further resolution to known population genetic structuring in the region, identifying the range-edge population in Maine as a distinct genetic cluster, and evidence of admixture with populations in New Brunswick and Quebec south of the St. Lawrence River. While previous research into trailing edge populations of Canada lynx have suggested that genetic diversity is lower than populations in the core of the distribution (Koen et al. 2014), our results show the opposite trend for Canada lynx in Maine. Relatively high genome-wide diversity, low levels of inbreeding, and evidence of admixture suggest that the peripheral Maine population may have high evolutionary potential powered by standing variation in neutral and adaptive loci. While the St. Lawrence River and Strait of Belle Isle strongly impede contiguous gene flow throughout the year; these barriers likely become at least temporarily permeable during the winter months when sea ice bridges enable lynx to migrate over the barrier (Koen et al. 2014). Koen et al. (2014) presented contemporary evidence of first-generation migration in both directions across the St. Lawrence River, including one individual that appeared to have traversed both the St. Lawrence and the Strait of Belle Isle. Despite our substantially reduced sample size, we similarly showed evidence of contemporary migration over the St. Lawrence River; with one individual from the NSLR cluster grouping with those from SSLR, and three individuals from SSLR and New Brunswick cluster grouping with the NSLR cluster. However, rising ambient and sea surface temperatures resulting from anthropogenic climate change could limit future opportunities for migration across sea ice and frozen stretches of water, potentially restricting ongoing gene flow.

Importantly, gene flow across the St. Lawrence River will hasten the core population’s adaptation to climate change through the adaptive introgression of warm-adapted loci from southern trailing-edge populations including Maine (Rehm et al. 2015). Contrary to other range edge populations of Canada lynx (Koen et al. 2014), individuals in Maine appear to have variable, yet relatively high levels of standing genomic variation, potentially indicating high evolutionary potential to future conditions. Although our analyses did not focus on detecting hybridization, the high levels of standing variation in the Maine population may be influenced by potential introgression of alleles from interspecies hybridization with bobcat (*Lynx rufus*: Homyack et al. 2008; Prentice et al. 2020), which are better adapted to warmer climates.

Genomic offset quantifies the magnitude of genetic change that would be required to maintain gene-environment relationships, which benefit individual fitness, under future conditions (Fitzpatrick and Keller 2015; Rellstab et al. 2021). In other words, the evolutionary potential required to evolve towards new physiological and behavioral optima under an “adapt in place” climate resilience strategy. An “adapt in place” resilience strategy is particularly relevant for island populations, such as Newfoundland. From a conservation perspective, mapping genomic offset can be a valuable tool for resiliency planning. Population-level offset can identify populations whose persistence will likely require on-the-ground conservation action to ensure the possibility of range-shift where evolutionary potential is low. Our results for the isolated population of lynx in Newfoundland reflect previous findings (Row et al. 2012; Koen et al. 2014). Pairwise comparisons identified the greatest genetic distances between Newfoundland and all other regions, with high levels of inbreeding and low levels of genome-wide diversity resulting from isolation. Our genomic vulnerability analyses show that western Newfoundland and the Gaspé Peninsula have the highest genomic offset over the coming century, suggesting little potential for an allelic shift to a new adaptive optimum under future conditions. Though the population is currently considered to be secure, these results accentuate concern for the future persistence of the Newfoundland population. Despite limited standing variation however, lynx on Newfoundland have rapidly evolved differences in body size, skull morphology, and diet consistent with Foster’s “island rule” (Khidas et al. 2013; Strong and Leroux 2014). Future work using coalescent analysis may further contextualize these results with insight into the colonization and demographic history of lynx in Newfoundland. Federal assessments by COSEWIC classify lynx as “not at risk” in Canada and lynx are legally harvested in 10 of 12 range provinces and territories, including the island of Newfoundland during an open season regulated by the Department of Fisheries, Forestry and Agriculture (Open Seasons Hunting and Trapping Order 2025). Management plans that strive to maintain or supplement standing variation by maintaining natural connectivity and/or introducing individuals from external populations could support the population’s long-term viability.

### Climate-associated adaptations in circadian entrainment and photoreception

Selective pressure shapes an organism’s receptivity to environmental cues, such as photoperiod, to ensure life-history events such reproduction are timed with conditions which maximize individual fitness (O’Malley et al. 2010). Standing variation in genes with strong gene-environment connections enables the optimal timing of life-history events to shift on pace with environmental change through rapid adaptation (Kondratova et al. 2010). Our results identified enrichment in ontology terms for circadian rhythm, circadian entrainment, and photoreception. NPAS2 is a core transcriptional activator which regulates circadian rhythm, by dimerizing with BMAL1 (DeBruyne et al. 2007) to regulate the master circadian clock in the hypothalamic suprachiasmatic nucleus (SCN) (Welsh et al. 2010, Reick et al. 2001). The regulation of circadian rhythm in the SCN is primarily driven through photoreception, with a suite of retinal circadian clock genes synchronizing cellular and molecular processes with the day/night light cycle. This link is important when considering the candidate adaptive loci presented here, which includes a neuropsin-producing gene (OPN5). Climate-associated OPN5 is expressed in intrinsically photosensitive retinal ganglion cells [ipRGCs]) ipRGCs, light-sensitive receptors which are provide light input to the SCN for photoentrainment (Yamazaki et al. 1999; Buhr et al. 2015). Previous studies have suggested that light is the most important Zeitgeber (signal) for entraining the mammalian circadian clock (Reppert and Weaver 2002; Sharma and Chandrasheckaran 2005; Golombek and Rosenstein 2010). For example, shortening photoperiod in autumn signals that temperatures are about to drop and the reverse occurs in spring. However, climate change affects the relationship between day length and temperature by causing temperatures to increase at a given latitude and by increasing the frequency of extreme weather events (IPCC 2013). Despite the impact on temperature and precipitation, global climate change has no effect on photoperiod (Andrews and Belknap 1993). Phenological mismatches between temperature and daylength have potential negative consequences for individual fitness (Stevenson et al. 2015; Walker et al. 2019). For example, the decrease in snow cover due to climate change can cause snowshoe hares to mis-time their seasonal color molt, a life event cued by photoperiod, exposing early moulting individuals to predation (Mills et al. 2013). Endogenous clocks must be plastic or rapidly adaptable in order to contribute to individual fitness under rapid environmental change (Bradshaw and Holsapfel 2010).

### Genetic load and Lynx susceptibility to idiopathic epilepsy

Genetic variants in a pool of standing variation can be advantageous, neutral (but potentially advantageous in the future) or disadvantageous. The “genetic load” of disadvantageous variation can incur fitness costs to reproduction and survival and is higher in small, inbred populations. Our assessment of genetic load in the 0.82 % of LD-pruned SNPs in the Canada lynx genome identified low and medium impact, largely synonymous mutations in genomic regions that contain genes linked to epilepsy and other seizure disorders. This included variation in GABRA1, an inhibitory neurotransmitter in the mammalian brain which, when pathogenically mutated, can cause idiopathic generalized epilepsy syndrome in juvenile humans (Cossette et al. 2002). Similarly, we identified synonymous variation in DSCAML1, which has been linked to seizure suppression (Hayase et al. 2020). Our analyses characterize the genomes of three lynx on Newfoundland as highly inbred, and mirror a susceptibility to epilepsy and seizure disorders documented in small, inbred populations of Iberian *(L. pardinus)* and Eurasian lynx *(L. lynx)* (Heaver and Waters 2019; Mínguez et al. 2021). For example, benign juvenile idiopathic epilepsy has been described in relatively high frequencies in kittens born within the Iberian lynx conservation breeding program (Heaver and Waters 2019; Kleinman-Ruiz et al. 2019; Mínguez et al. 2019; Mínguez et al. 2021). To our knowledge, there are no reports of Canada lynx with clinical signs of epilepsy or other seizure disorders in Newfoundland. Our findings present a candidate list of genes for exploring the causative genetic basis of idiopathic epilepsy in *Lynx* species.

### Implications for conservation management at the trailing edge of the Canada lynx distribution

This study is the first direct assessment of evolutionary potential and genetic offset for Canada lynx populations and provides important baseline assessments for strategic recovery planning. Canada lynx within the contiguous United States have been listed as threatened under the Endangered Species Act (ESA) since 2000. In 2018, the U.S. Fish and Wildlife Service (USFWS), the regulating authority for Canada lynx in the contiguous United States, withdrew a previously proposed rule to delist the species, concluding that available evidence did not support delisting at that time (USFWS 2018). The species’ status was subsequently re-evaluated in a five-year review completed in 2022, which again retained the Canada lynx as a threatened species under the Endangered Species Act and outlined a 2024 recovery plan in which climate change was the leading threat to the long-term persistence of populations in the contiguous United States (USFWS 2022, USFWS 2024). Our results suggest that the population of Canada lynx in Maine is rich in neutral and climate-adapted variation. If gene flow allows for adaptive introgression, genomic variation in the Maine population may help regional populations adapt to the rapidly changing climate.

## Supplementary Material

See attached Supplementary Tables and Figures

## Declarations

TL conceived the study. TL, JV, LMK, JK, KL, KB, JB, PW participated in sampling and/or laboratory work. TL, BB performed bioinformatics, population genomics, identification of outlier loci, genotype-environment association tests, functional annotations, and gene set enrichment tests. TL, BB performed selection and analysis of climatic covariables. BB parameterized, curated, and documented analysis pipelines. SDS, WJ, JO, JV provided support with initial study planning and interpretation of results. TL, BB drafted the manuscript with substantial input from LMK, WJ, and JO. All authors read and approved the final manuscript.

Code used for all analyses can be accessed at https://github.com/bpbentley/lynx_wgr. The authors claim no conflicts of interest.

## Supporting information

SupTables

SupFigures

## Acknowledgements

The authors thank Drs. Jeff Bowman, Paul Wilson and Melanie Prentice for generously providing samples for this effort. Dr. Brenna Forester provided guidance on genotype-environment association methods. The authors acknowledge the wildlife photography of Bill Byrne (MassWildlife) featured in Figure 1. This work was funded by the U.S. Fish and Wildlife Service and the Maine Department of Inland Fisheries & Wildlife Grant application for Pittman-Robertson Wildlife Restoration Act federal assistance #W-86-R and utilized resources from Unity, a collaborative, multi-institutional high-performance computing cluster managed by UMass Amherst Research Computing and Data. Co-first authors TL and BB contributed equally to this manuscript and each has the right to list themselves first in author order on curricula vitae.

