## Supplementary material for "Assessing climate adaptation among Canada lynx (*Lynx canadensis*) populations at the trailing edge": SupFigures

**Supplementary Figures**

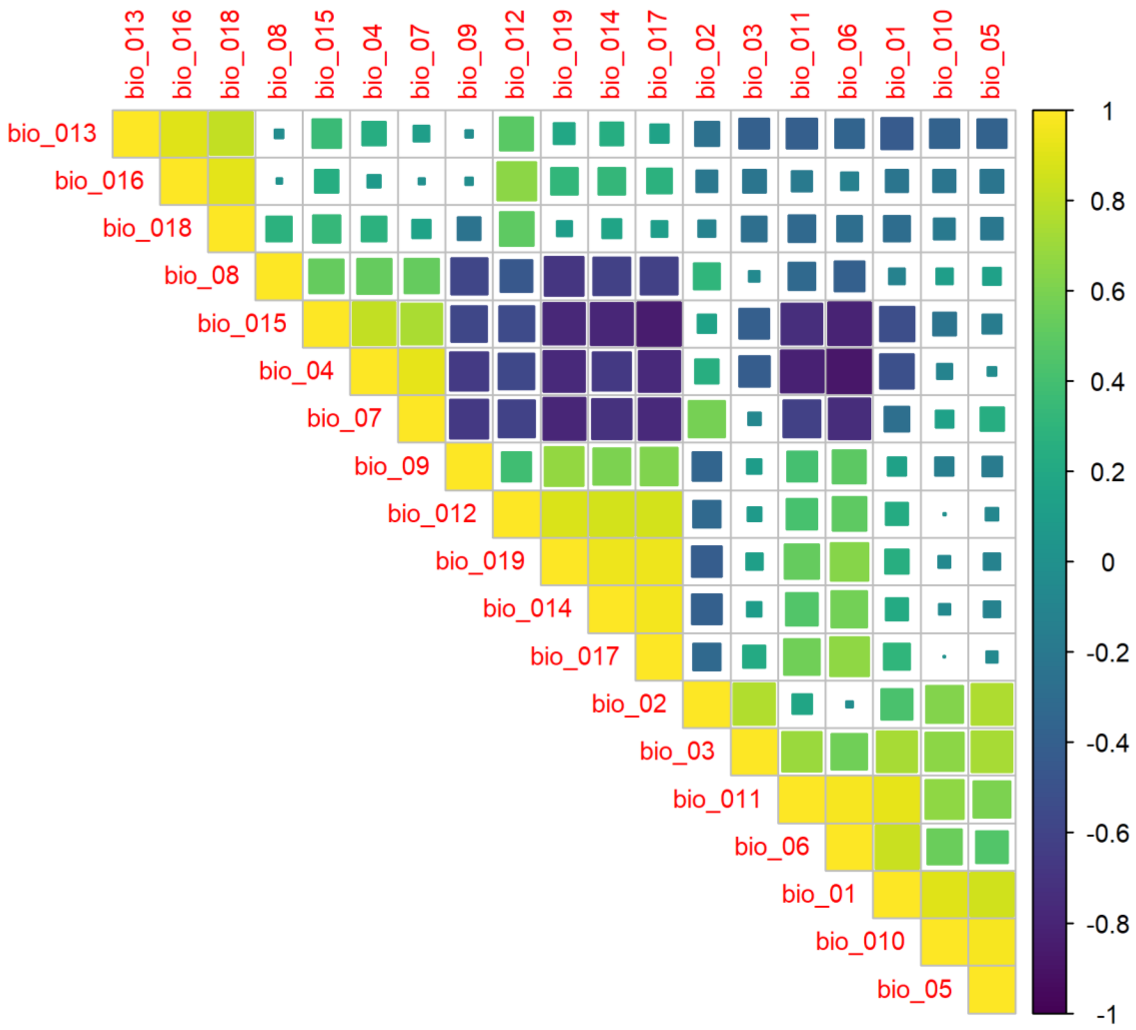


**Figure S1.** Pairwise correlations between Bioclimatic variables extracted from the WorldClim v2.1 database. Variables were selected such that no correlations of R >0.8 were retained in the final dataset (n=8).

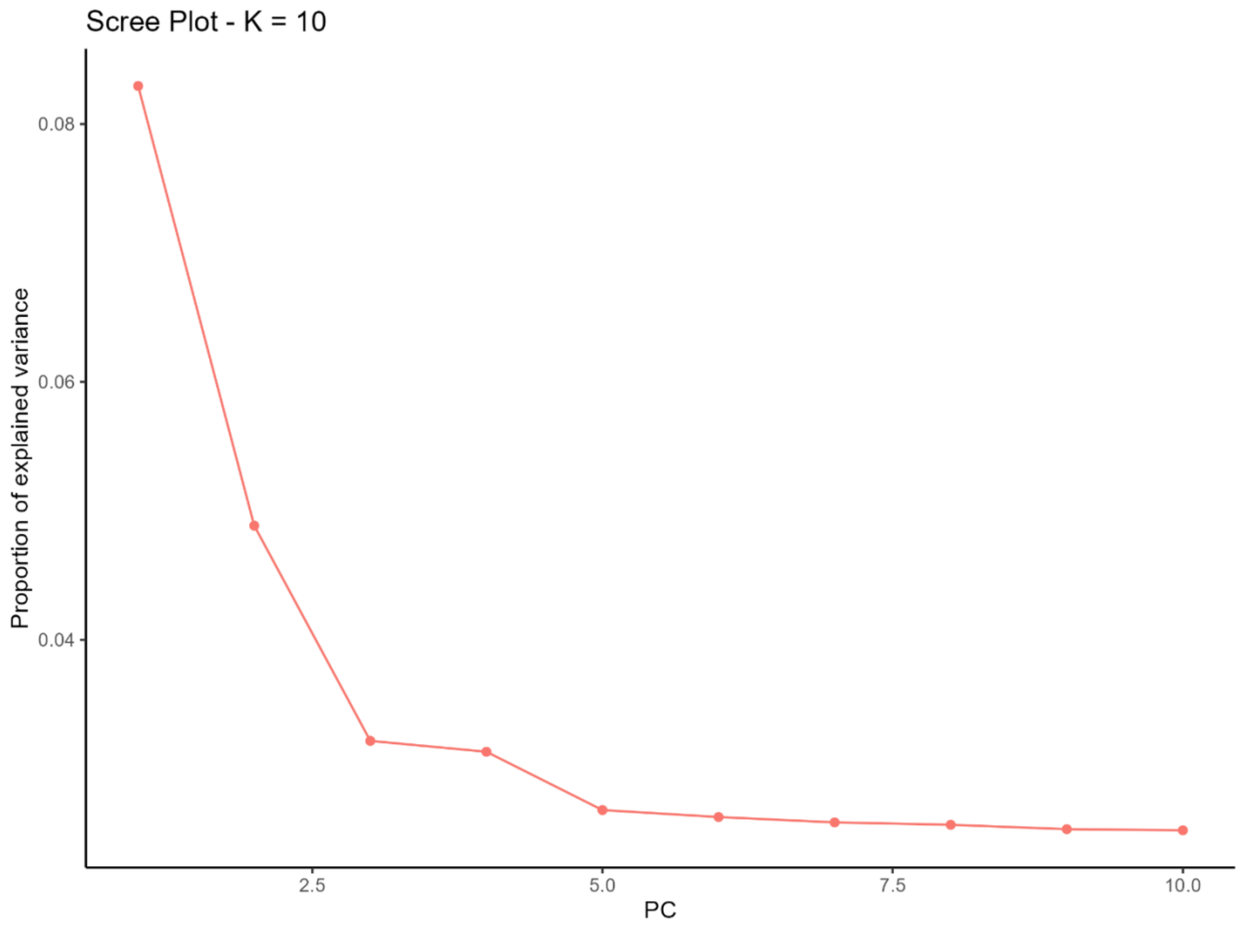


**Figure S2.** Scree plot with the first 10 clusters (K) calculated from PCAdapt. Using the elbow-method, K=4 was chosen for downstream analyses.

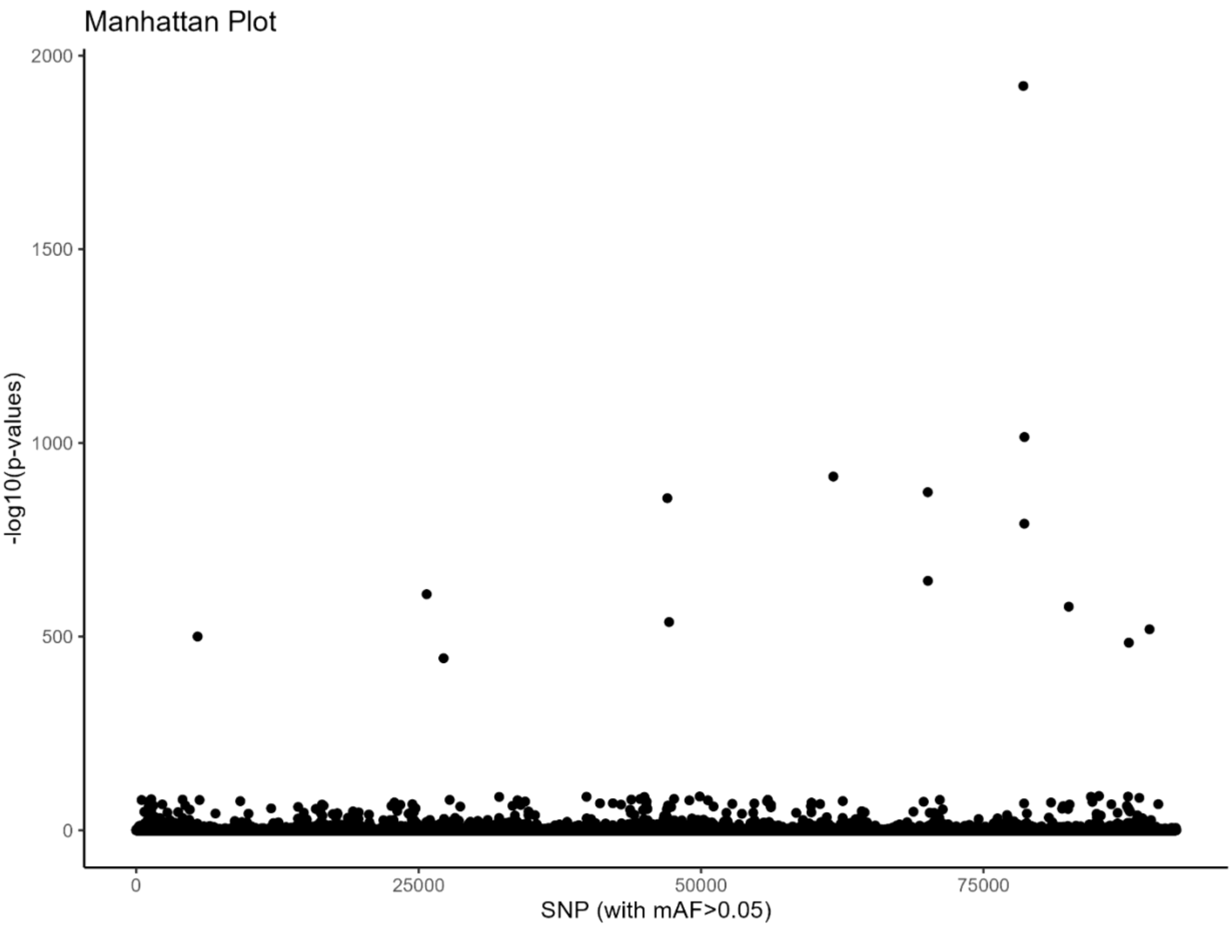


**Figure S3.** Manhattan plot of outliers identified by PCAdapt with K=4.

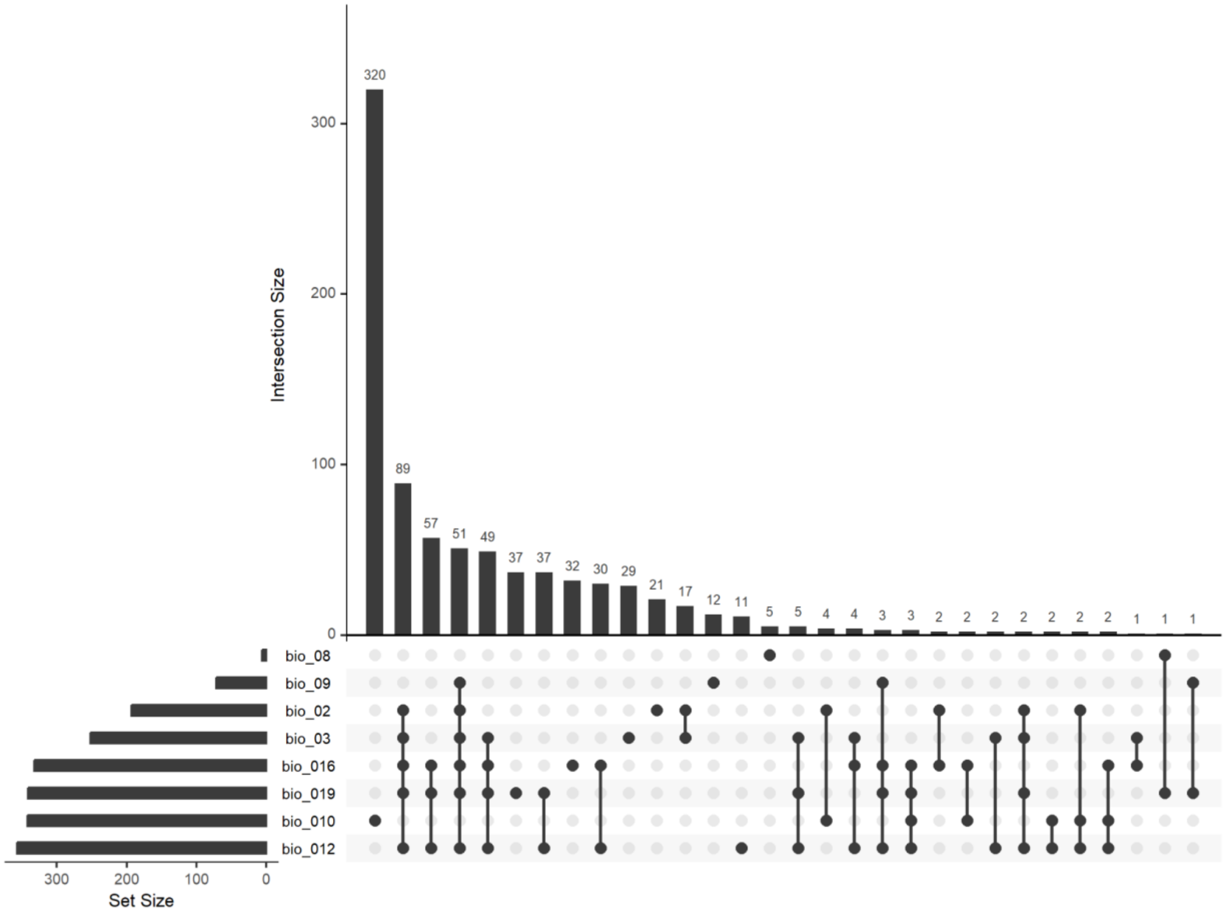


**Figure S4.** Upset plot showing the relationship between loci identified through the LFMM univariate analysis, and each of the bioclimatic variables.

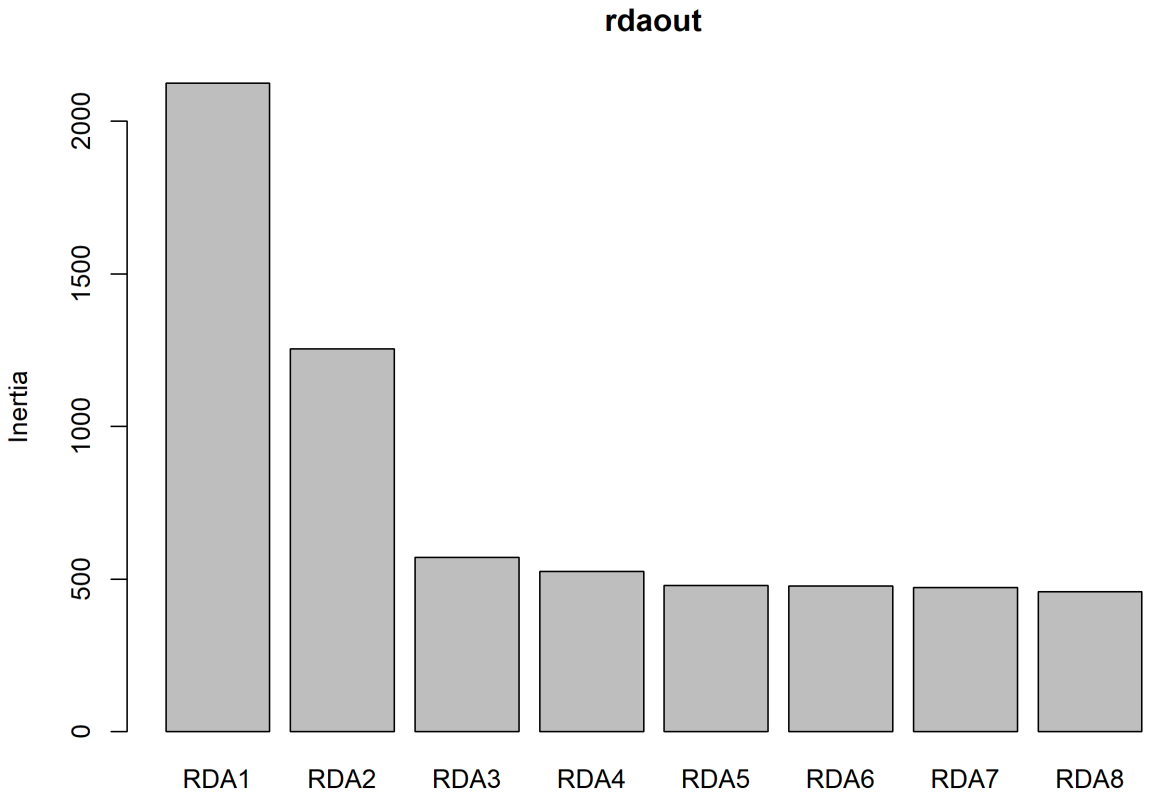


**Figure S5**. Scree plot from redundancy analysis (RDA) showing the variance (inertia) explained by each canonical axis. The first two axes (RDA1 and RDA2) explain the majority of the constrained genetic variation, indicating strong multivariate associations with environmental variables.

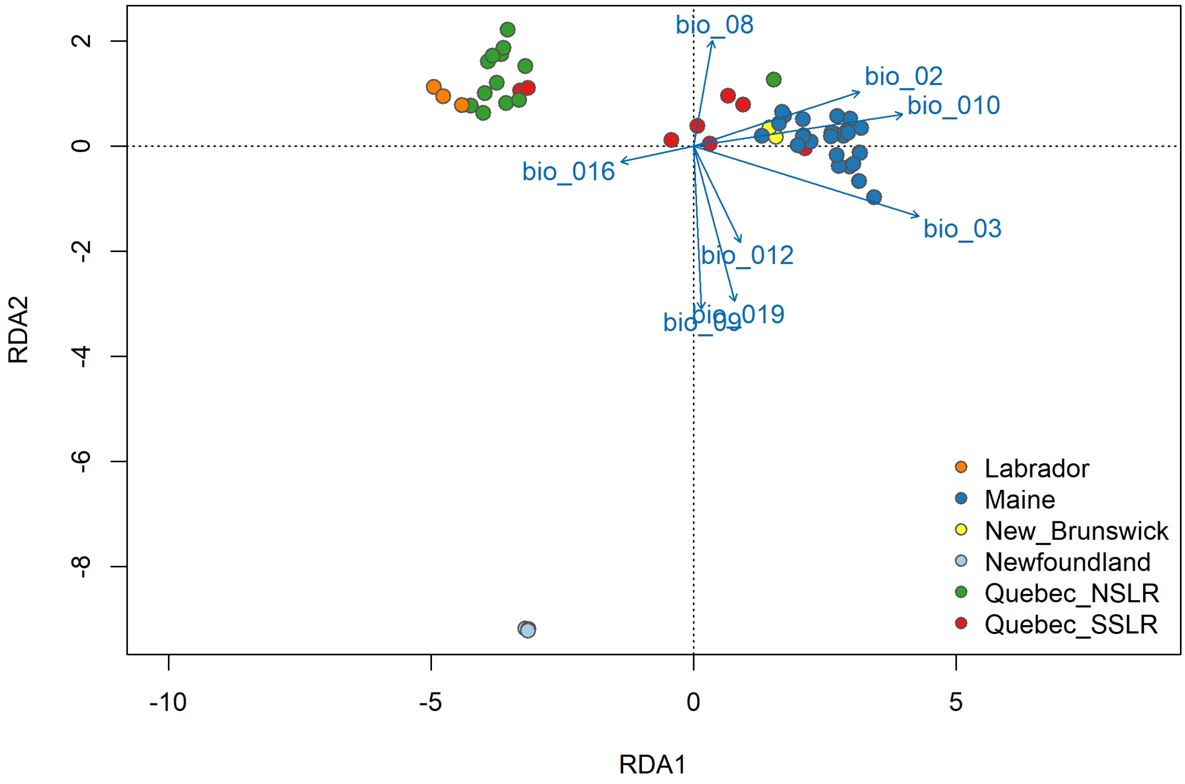


**Figure S6**. Redundancy analysis (RDA) biplot showing the relationship between individual genotypes and environmental variables across six geographic populations of *Lynx canadensis*. Colored points represent individuals from different collection regions, and blue arrows indicate the direction and strength of the association between standardized BioClim variables and genetic variation. The first two RDA axes explain 53.1% of the constrained genetic variance.

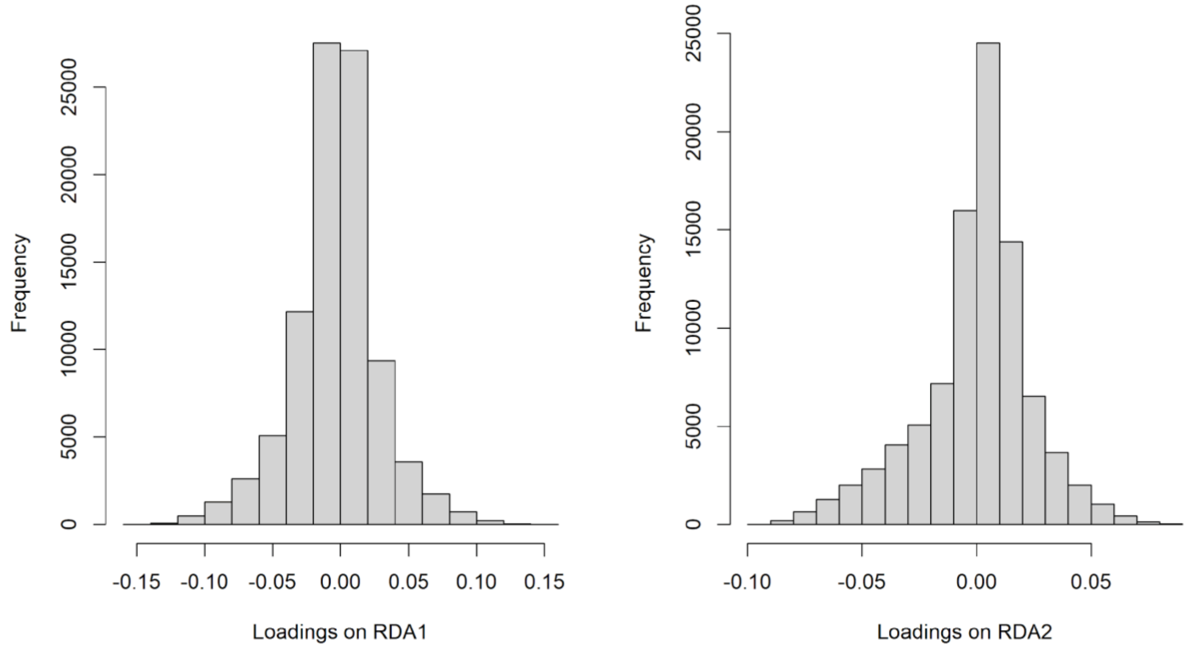


**Figure S7**. Loadings of each locus on the two retained RDA axes.

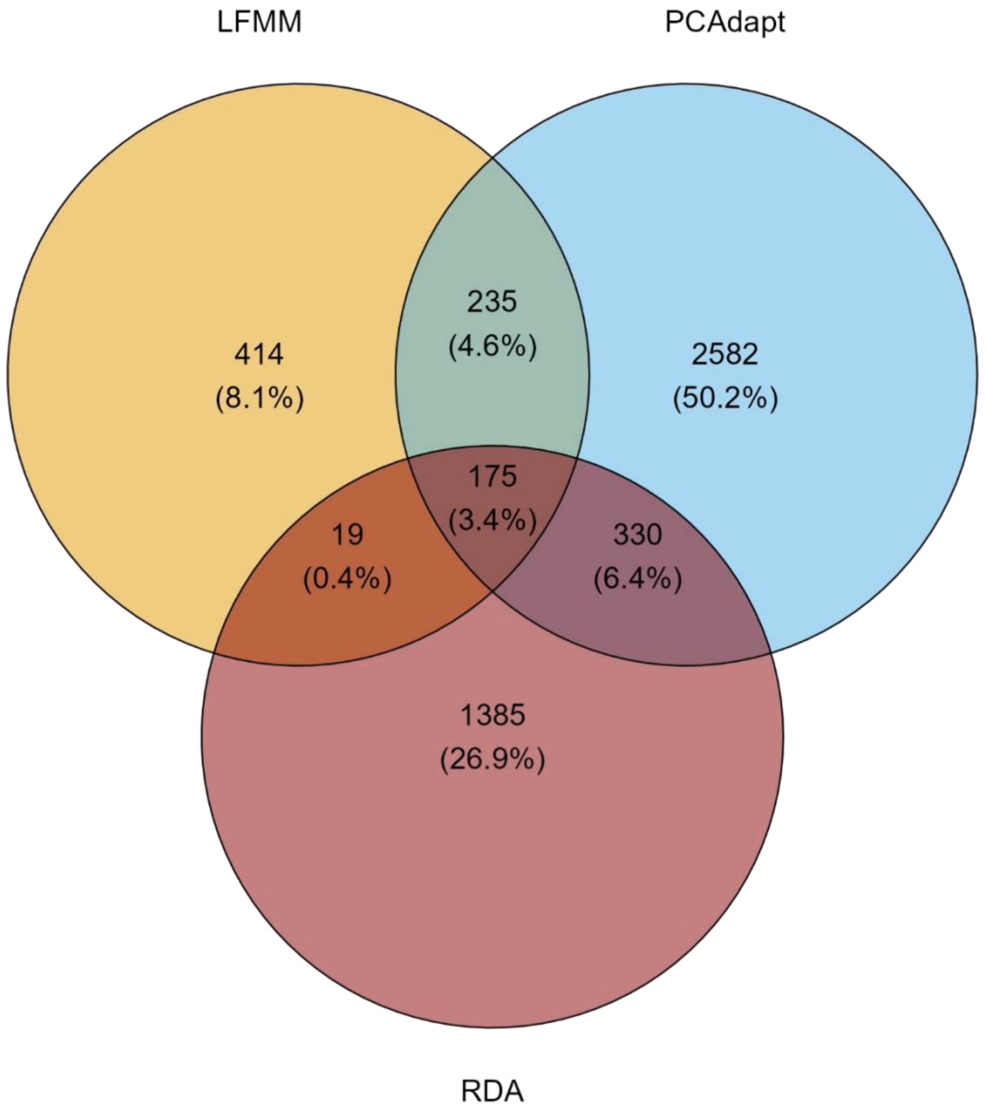


**Figure S8.** Total number, and number of overlapping loci identified by the three independent outlier detection tests. Latent factor mixed-models (LFMM) and redundancy analyses (RDA) associate loci with specific bioclimatic variables (genotype-environment association analyses; GEA), while PCAdapt does not.
